# Genic evidence that gnetophytes are sister to all other seed plants

**DOI:** 10.1101/629915

**Authors:** Yinzhi Zhang, Zhiming Liu

## Abstract

Gnetophytes, comprising three relict genera, *Gnetum*, *Welwitchia* and *Ephedra*, are a morphologically diverse and enigmatic assemblage among seed plants. Despite recent progress on phylogenomic analyses or the insights from the recently decoded *Gnetum* genome, the relationship between gnetophytes and other seed plant lineages is still one of the outstanding, unresolved questions in plant sciences. Here, we showed that phylogenetic studies from nuclear genes support the hypothesis that places gnetophytes as sister to all other extant seed plants and so this hypothesis should not be ruled out according to phylogenetic inference based on nuclear genes. However, this extraordinarily difficult phylogenetic problem might never be solved by phylogenetic inference based gene tree under various artificial selection. Hence, we adopted a novel approach, comparing gene divergence among different lineages, to solve the conflicts by showing that gnetophytes actually did not gained a set of genes like the most recent common ancestor (MRCA) of other seed plants. This distinct gene evolution pattern could not be explained by random gene lost as in other seed plants but should be interpreted by the early divergence of gnetophytes from rest of seed plants. With such a placement, the gymnosperms are paraphyletic and there should be three distinct groups of living seed plants: gnetophytes, non-gnetophytes gymnosperms and angiosperms.

## Introduction

Although the living seed plants exhibit a high degree of species richness and extensive morphological variation, extant seed plant taxa represent only a relic of their former diversity. The extensive extinction and long independent evolution and convergence in both morphological characters and genome sequences, has made it extremely difficult to correctly trace phylogenetic relationships among seed plant lineages. This problem has been triggering debates for over several decades and the central issue is the placement of an enigmatic group of gymnosperms called the genophtes (Fig. 1): The ‘anthophyte hypothesis’ was first proposed because of shared morphological similarities with angiosperms. This hypothesis placed gnetophytes as most closely related to angiosperms (Crane, 1985; Doyle & Donoghue, 1986). However, this hypothesis had been ruled out due to the rise and development of molecular research (Doyle et al., 1994; Wickett et al., 2014). Most molecular studies based on plastid and nuclear sequence have show strong support to the hypothesis that gnetophytes should be placed closely related to conifers. It mainly includes three hypotheses: either as sister to Pinaceae (‘Gnepine’), to cupressophytes (‘Gnecup’), or to all conifers (‘Gnetifer’), depending on the different data sets, sites selection, and inferring methods involved (Ruhfel et al., 2014; Wickett et al., 2014). Moreover, sophisticated phylogenomic analyses based on large-scale data matrices are also pointing towards the hypothesis that gnetophytes are most closely related to conifers (Wickett et al., 2014; Ran et al., 2018). Interestingly, the other two remaining hypotheses, that have received little attention, but have also been constantly supported using different sets of gene sequence and various phylogenomic reconstruction methods, either place gnetophytes as sister to all other gymnosperms (Sanderson et al., 2000; Rai et al., 2008; Li et al., 2017) or even all seed plants (Frohlich & Parker, 2000; Schmidt & Schneider-Poetsch, 2002; Lee et al., 2011; Chen et al., 2016). Notably, the ‘Seed plants sister’ hypothesis has not been supported with phylogenetic tree inferred from nuclear genes to our current knowledge and tends to be ignored in phylogenetic testing (Wickett et al., 2014).

**Figure 1.**
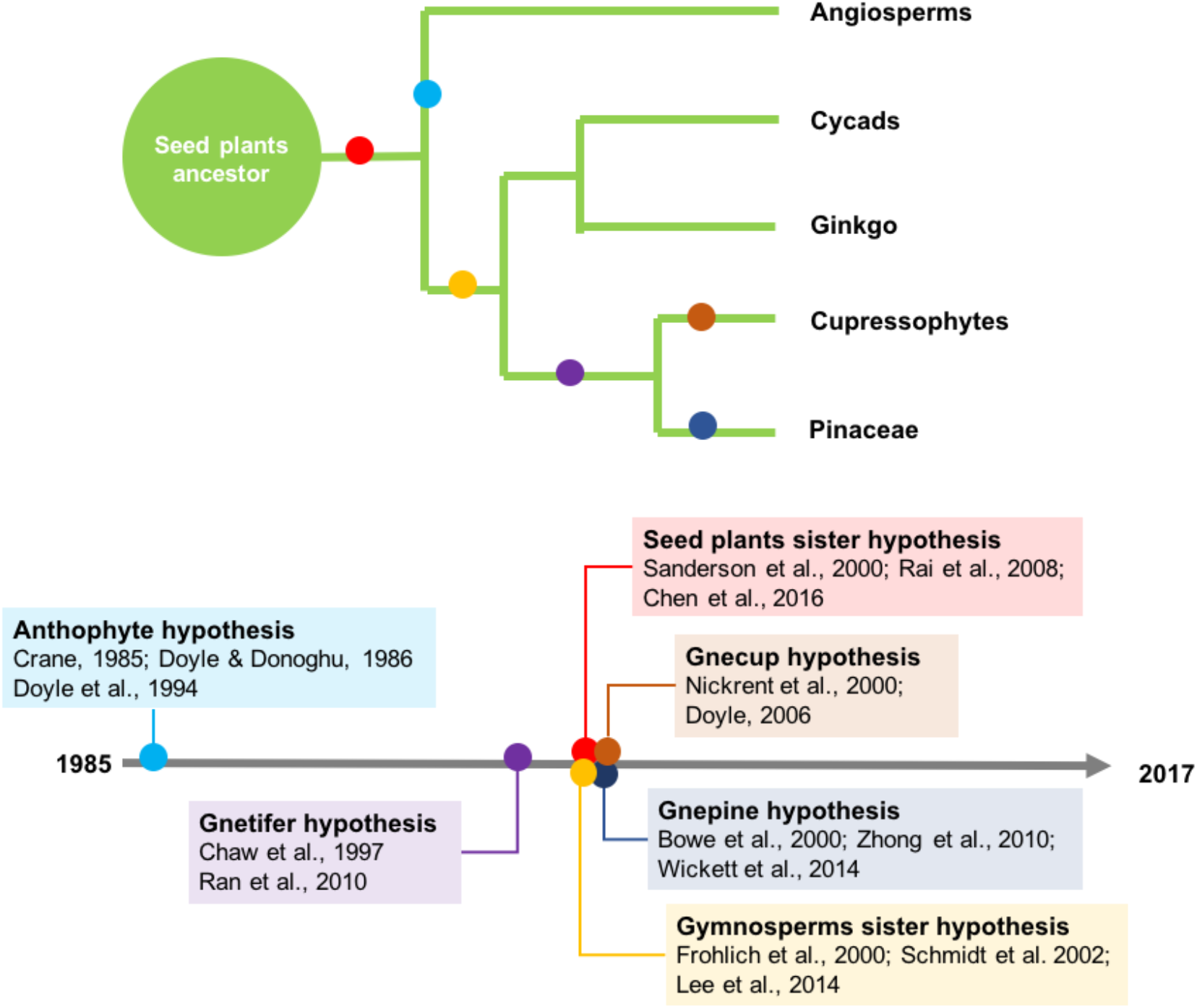
Six conflicting hypotheses of the phylogenetic position of gnetophytes inferred by diverse sets of molecular markers or morphological character clustering. The anthophytes hypothesis was first proposed due to morphological character cladistic and once triggered great debates on history. Couple of genes or alignments from large matrices of nuclear or plastid genome compartments were conducted and provide conflicting topologies on phylogenetic positions of gnetophytes.

### Phylogenetic inferences on nuclear genes

To test the strength of the emerging consensus hypothesis that gnetophytes are closed to conifers, we measured the phylogenetic signals from a data matrix of 13 land plants species (using nine whole genome assemblies and four whole transcriptome data sets, Table 1). We used the whole genome assembly of *G. montanum*, and transcriptomic data from *Ephedra equisetina* (Wan et al., 2018, see ‘*Gnetophytes Genome Project*’, which also contains information on data production, taxon sampling, assembly and annotation). We also used genomic data from nine published plant genomes and transcriptome data of Cycads and *Ginkgo* (Table 1; six gymnosperms - *G. montanum*, *E. equisetina*, *Picea abies*, *P. taeda*, *G. biloba*, and *Cycas elongata*; six angiosperms - *A. trichopoda*, *Vitis vinifera*, *Arabidopsis thaliana*, *Musa acuminata*, *Elaeis guineensis*, *and Oryza sativa*, and; one non-seed plant - *S. moellendorffii*). Protein genes of species with genomes were from database (Table 1). For species with transcriptomic data, we used Genewise 2.4.1 (Birney et al., 2004) to generate gene structures based on proteins to the assembled *G. montanum* sequence. Protein gene from all these species were used for phylogenetic analyses.

**Table 1.** Species and genome assemblies used in this study

| Species | Sources of data |
| --- | --- |
| *Selaginella moellendorffii* | http://genome.jgi.doe.gov/pages/dynamicOrganismDownload.jsf?organism=PhytozomeV10 |
| *Picea abies* | http://congenie.org |
| *Pinus taeda* | http://pinegenome.org/pinerefseq/ |
| *Ginkgo biloba* | [ftp://ftp.plantbiology.msu.edu/pub/data/MPGR/Ginkgo biloba/](ftp://ftp.plantbiology.msu.edu/pub/data/MPGR/Ginkgo_biloba/) (EST data) |
| *Cycas elongata* | http://journals.plos.org/plosone/article?id=10.1371/journal.pone.0154384 |
| *Gnetum montanum* | https://www.ncbi.nlm.nih.gov/bioproject/PRJNA339497 |
| *Ephedra equisetina* | https://www.ncbi.nlm.nih.gov/bioproject/PRJNA339497 |
| *Amborella trichopoda* | ftp://ftp.ensemblgenomes.org/pub/plants/release-25/plants/ |
| *Musa acuminata* | ftp://ftp.ensemblgenomes.org/pub/release-25/plants/ |
| *Oryza sativa* | ftp://ftp.ensemblgenomes.org/pub/release-25/plants/ |
| *Vitis vinifera* | ftp://ftp.ensemblgenomes.org/pub/release-25/plants/ |
| *Arabidopsis thaliana* | ftp://ftp.ensemblgenomes.org/pub/release-25/plants/ |
| *Elaeis guineensis* | ftp://ftp.ensemblgenomes.org/pub/release-25/plants/ |

Similarities between these proteins were detected using an all-against-all BLASTp version 2.2.26 (Altschul et al., 1997) with an E-value of 1e^−10^. Only alignments between gene pairs that had >0.5 coverage of each sequence were retained for analysis. The software OrthoMCL (Li et al., 2003), based on a Markov cluster algorithm was applied to find orthogroups (with an inflation value of 1.5, to extract orthologous and paralagous proteins). Then we retrieved a single copy sequence in per species, to generate a 1,334 gene dataset. Then the alignments of 331,467 AA sites were concatenated to a super alignment matrix. ProtTestwas used to select the best fit model (LG+Γ4 model) for amino acid replacement and RA×ML (v8.0.19) was used to reconstruct a maximum likelihood tree. Robustness of the maximum likelihood tree was assessed using the bootstrap method (100 pseudo-replicates). A ML tree supporting “gnetophytes around the conifers” were shown in Fig. 2A.

**Figure 2.**
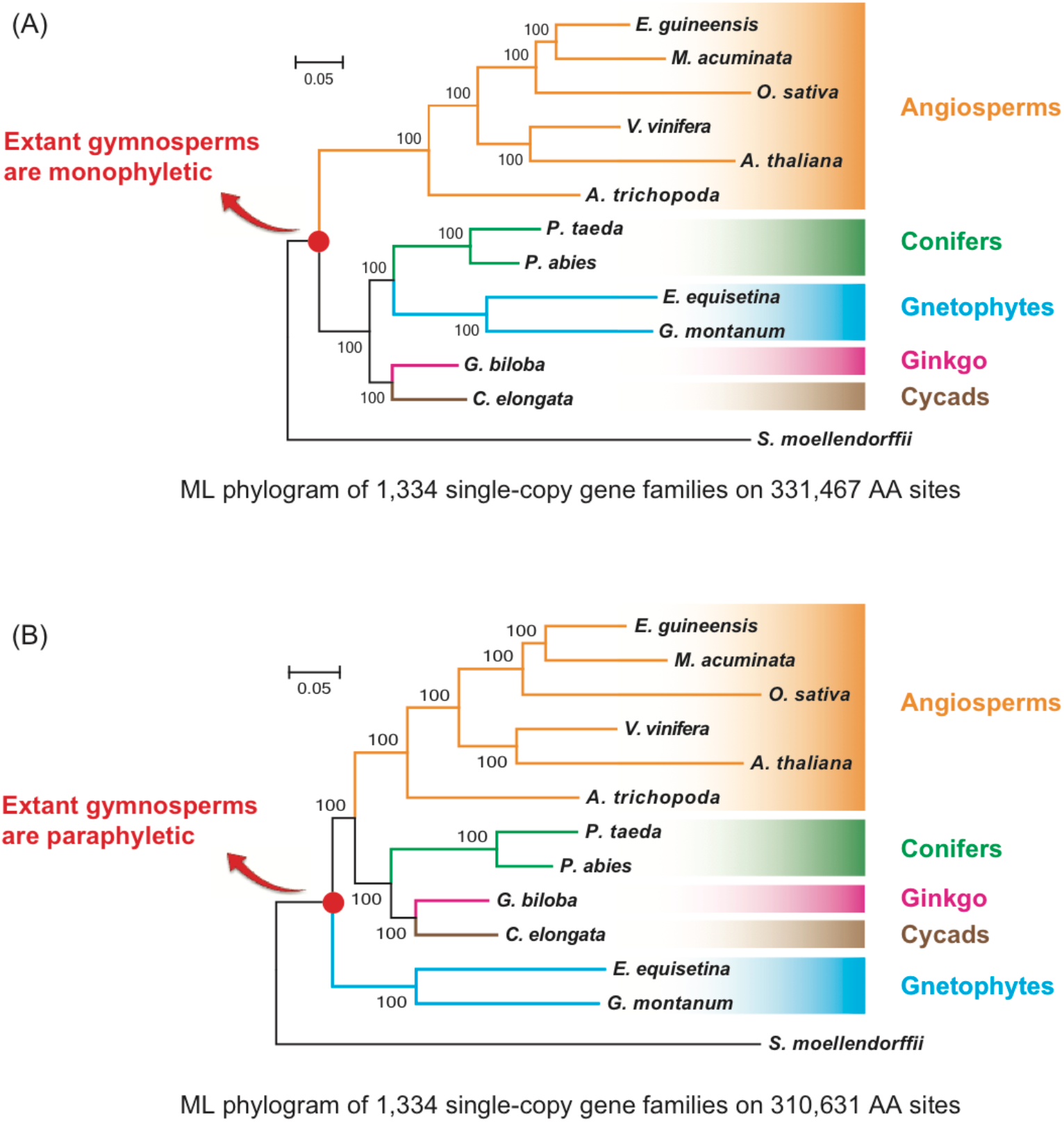
Phylogenetic inference based on nuclear genes with different sites selection. The alignments of 331,467 AA sites in Figure 2A (or 310,631 AA sites in Figure 2B) were concatenated to a super alignment matrix. ProtTest was used to select the best fit model (LG+Γ4 model) for amino acid replacement and RA×ML (v8.0.19) was used to reconstruct a maximum likelihood tree. Robustness of the maximum likelihood tree was assessed using the bootstrap method (100 pseudo-replicates).

Considering the following three factors: (a) The extinctions of the ancestor of modern angiosperms; (b) angiosperms have greater average rate of substitution than gymnosperms; (c) the slow evolutionary rate of the gymnosperms, the distinct evolution rates among extent angiosperms and gymnosperms as well as the lack of ancient angiosperm ancestor sequence, simultaneously leads to that gnetophytes will contain much more same amino-acids with the non-gnetophyte gymnosperms than extant angiosperms, which will outweigh the real evolutionary signals during phylogenetic inference. Hence, to reduce the effect of (a) and (b), we retained sites where at least one of the six gymnosperms has same amino-acid with at least one of the six angiosperms. Finally, 310,631 (94%) of the total 331,467 sites were retained and used to do the phylogenic tree construction following the above approach. Notably, a ML tree (with 100% bootstrap support) indicating “gnetophytes as sisters to other seed plants” based on these sites were generated and showed in Fig. 2B.

Though the ‘gnetophytes as sister to other seed plants’ hypothesis has been reported previously, either by using whole plastid sequence data, plastid proteins, or mitochondrial genes (data were not shown) analyzed with a range of methods including maximum parsimony and maximum likelihood (Frohlich & Parker, 2000; Schmidt & Schneider-Poetsch, 2002; Lee et al., 2011; Chen et al., 2016), this hypothesis tends to be ruled out based on nuclear loci (Wickett et al., 2014). Here, for the first time, we showed that this hypothesis could also be supported from phylogenetic inference from nuclear genes. Given this, a hypothesis of gnetophytes being sister to all other seed plants cannot be ruled out on the basis of phylogenetic trees inferred from nuclear loci. However, resolving phylogenetic relationships among extant seed plants has been shown as an extraordinarily difficult problem only based on gene sequences (Wickett et al., 2014). Hence, we think there is little hope that the mainstream approach of gene tree construction resulting from different treatments of the data and methods of analysis will solve this problem fundamentally. We should adopt other novel approach to provide better resolution of relationships among major seed plant clades.

### Gene evolution patterns among representative seed plants

Because of the inconsistencies in signals using typical phylogenetic approaches, we developed an indirect approach to examine patterns of gene proliferation among major seed plant clades. In total, five different protein coding gene sets in *Amborella trichopoda* were compared against the relevant protein coding genes in *Sellaginella moellendorffii* (Fig. 3, supplementary table 1-5). Only genes of *A. trichopoda* having at least a score > 50 with *S. moellendorffii* were used in the boxplot. The determination and selection methods are as follows:

**Figure 3.**
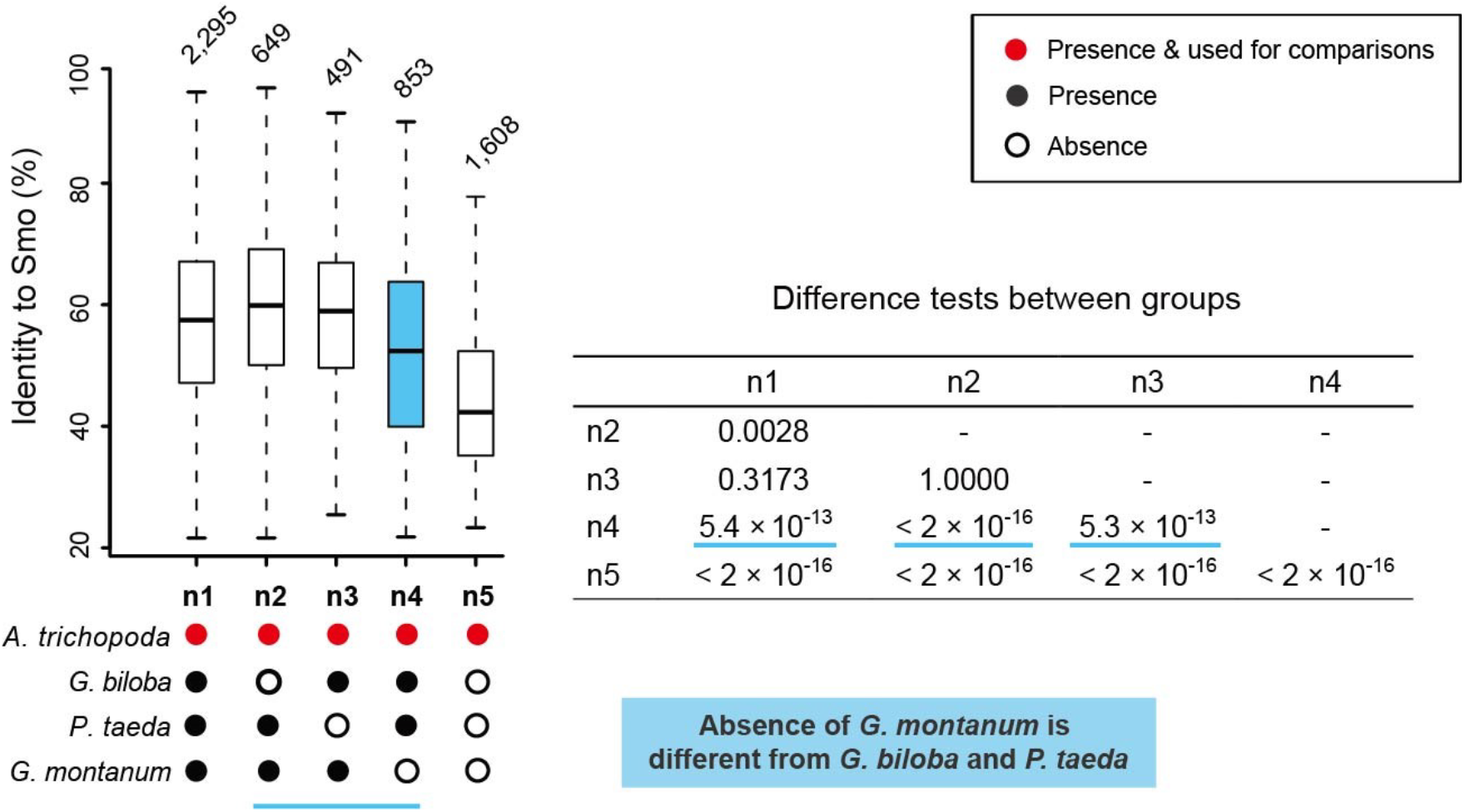
Genic signatures for gnetophytes being distinct from other seed plants. Comparisons of *A. trichopoda* gene sets that carry orthologues present in different combinations of gymnosperms (BLASTp gene identities against *S. moellendorffii*-Smo). The “n1” group contains genes found in all seed plants investigated and were likely found in the seed plant ancestor, while “n5” represents genes present only in *A. trichopoda*, genes that are likely to have diverged substantially only in the angiosperm lineage). When *G. montanum* orthologous genes are absent (“n4”), the dataset has significantly lower overall BLASTp identities than in datasets where orthologues are absent in other gymnosperms (e.g. “n2” and “n3”).

a. We defined an “n1” set of single copy orthoMCL genes (Li et al., 2003), which are those groups shared by *G. montanum*, *G. biloba*, *P. taeda* and *A. trichopoda* (supplementary table S2).
b. The “n2” gene set comprised protein-coding genes considered to be missing in *G. biloba* but present in the other seed plants, *G. montanum*, *P. taeda* and *A. trichopoda.* We used BLASTp, with an E-value threshold of 1e^−5^, to query protein sets from *G. montanum* and *P. taeda* against protein sets from *G. biloba* and *A. trichopoda*. Genes shared by *G. montanum* and *P. taeda*, that also had ≥ 10% higher scores (score=identity*coverage) when queried against *A. trichopoda* compared with *G. biloba*, were considered to be genes that are absent in *G. biloba*.
c. The “n3” set were protein genes considered to be missing in *P. taeda* but present in the other seed plants, *G. montanum*, *G. biloba*, *A. trichopoda.* We aligned by BLASTp with an E-value of 1e^−5^ protein sets from *G. biloba* and *G. montanum* against protein sets from *P. taeda* and *A. trichopoda*. Genes shared by *G. biloba* and *G. montanum*, that also had ≥ 10% higher scores (score=identity * coverage) when aligned against *A. trichopoda* compared with *P. taeda*, were considered to be genes that are absent in *P. taeda*.
d. The “n4” set consisted of protein genes considered to be missing in *G. montanum* but present in the other seed plants, *G. biloba*, *P. taeda*, *A. trichopoda.* We aligned by BLASTp with an E-value of 1e^−5^ protein sets from *G. biloba* and *P. taeda* against protein sets from *G. montanum* and *A. trichopoda*. Genes shared by *G. biloba* and *P. taeda*, that also had ≥ 10% higher scores (score=identity*coverage) when aligned against *A. trichopoda* compared with *G. montanum*, were considered to be genes that are absent in *G. montanum*.
e. The “n5” set were protein genes considered to occur only in *A. trichopoda* specific families.

We mainly focused on genes that were present in *Amborella trichopoda* but were absent in one of the gymnosperms (i.e. absent in either *G. montanum*, or *Ginkgo biloba*, or *Pinus taeda*). The “n1” set, with a speculative orthologue in all the seed plant lineages, were considered to be genes that have been retained from the seed plant ancestor (Supplementary Table 1). In contrast the “n2”, “n3” and “n4” sets have an orthologue missing in *G. biloba*, *P. taeda* or *G. montanum* (Supplementary Table 2-4). The “n5” set of genes (Supplementary Table 5) is considered to be specific to *A. trichopoda* or gained after angiosperms separated from gymnosperms. Our analyses of the sequence identities of the “n1”-“n5” gene sets showed that the sequence identity distribution of the “n5” set is lower than the “n1” set (*p* < 2 × 10^−16^, Fig. 3). Remarkably, the “n4” set, representing genes thought to be absent in *G. montanum*, also has a lower sequence identity distribution than that of the “n2” set (orthologous genes considered to be absent in *G. biloba*, *p* < 2 × 10^−16^, Fig. 3) and the “n3” set (orthologous genes considered to be absent in *P. taeda*, *p* < 5.3 × 10^−13^, Fig. 3). When *G. montanum* orthologous genes are absent (“n4”), the dataset has significantly lower overall BLASTp identities than in datasets where orthologues are absent in other gymnosperms (e.g. “n2” and “n3”). What becomes apparent is that orthologous genes in *G. montanum* (i.e. n1, n2, and n3 sets; Fig. 3) play a notable role in comparison among all gene sets identity distributions (Fig. 3).

How to interpret this gene evolution patterns? It’s important to note that these analyses are not affected by the different substitution rate among *G. montanum*, *G. biloba*, and *P. taeda* since all the genes of “n1”-“n5” sets are from *A. trichopoda*. Firstly, “n5” set is deservedly considered later obtained by *A. trichopoda* after its split with other gymnosperms and the genes from “n5” set are more young than other genes from other sets (Fig. 4). Hence, the “n5” set genes has less sequence similarities (or identities) with the ancient *S. moellendorffii* genes. It is expected that because comparison between orthologs of ancient, conserved (“old”) genes would be more similar to each other, and have higher sequence identities than orthologs of less conserved, or more rapidly diverging (“yound”) genes. Secondly, “n2” genes and “n3” genes show no obvious difference, which indicated that “n2” genes and “n3” genes were obtained before the split of *G. montanum*, *G. biloba*, *P. taeda* and *A. trichopoda* (in other words, these genes were obtained in the common ancestor of *G. montanum*, *G. biloba*, *P. taeda* and *A. trichopoda*). The absence of these genes in *G. biloba* or *P. taeda* is due to random lost during their independent evolution (Fig. 4). And this speculation is confirmed by that there are also no obvious difference between “n1” genes and “n2”/“n3” genes since “n1” represents genes shared by *G. montanum*, *G. biloba*, *P. taeda* and *A. trichopoda* (in other words, these genes were also obtained in the common ancestor of *G. montanum*, *G. biloba*, *P. taeda* and *A. trichopoda*). In other words, “n2”/“n3” genes were as “old” as “n1” genes. Notably, the ‘gene absence’ patterns could not be explained by random lost in *G. montanum* but should be interpreted by the early divergence of gnetophytes from rest of seed plants because “n4” has much lower identities than “n1”, “n2”, and “n3”. In other words, part (red G2 in Fig. 4) of “n4” genes were actually obtained only by the common ancestor of *G. biloba*, *P. taeda* and *A. trichopoda* with early split of *G. montanum* and these “n4” genes are “younger” than “n1”, “n2”, and “n4” but “older” than “n5” genes. The observed gene evolution pattern is only consistent with the hypothesis that gnetophytes are sister to all other extant seed plants (Fig. 4). Notably, that there is little overlap (Wayne comparison in Fig. 4) between these three sets (‘n2’, ‘n3’ and ‘n4’) also confirms than our approach can detect specific gene absence in each species, which are not affected by the gene loss in other nodes (such as L4 or L2 in Fig. 4).

**Figure 4.**
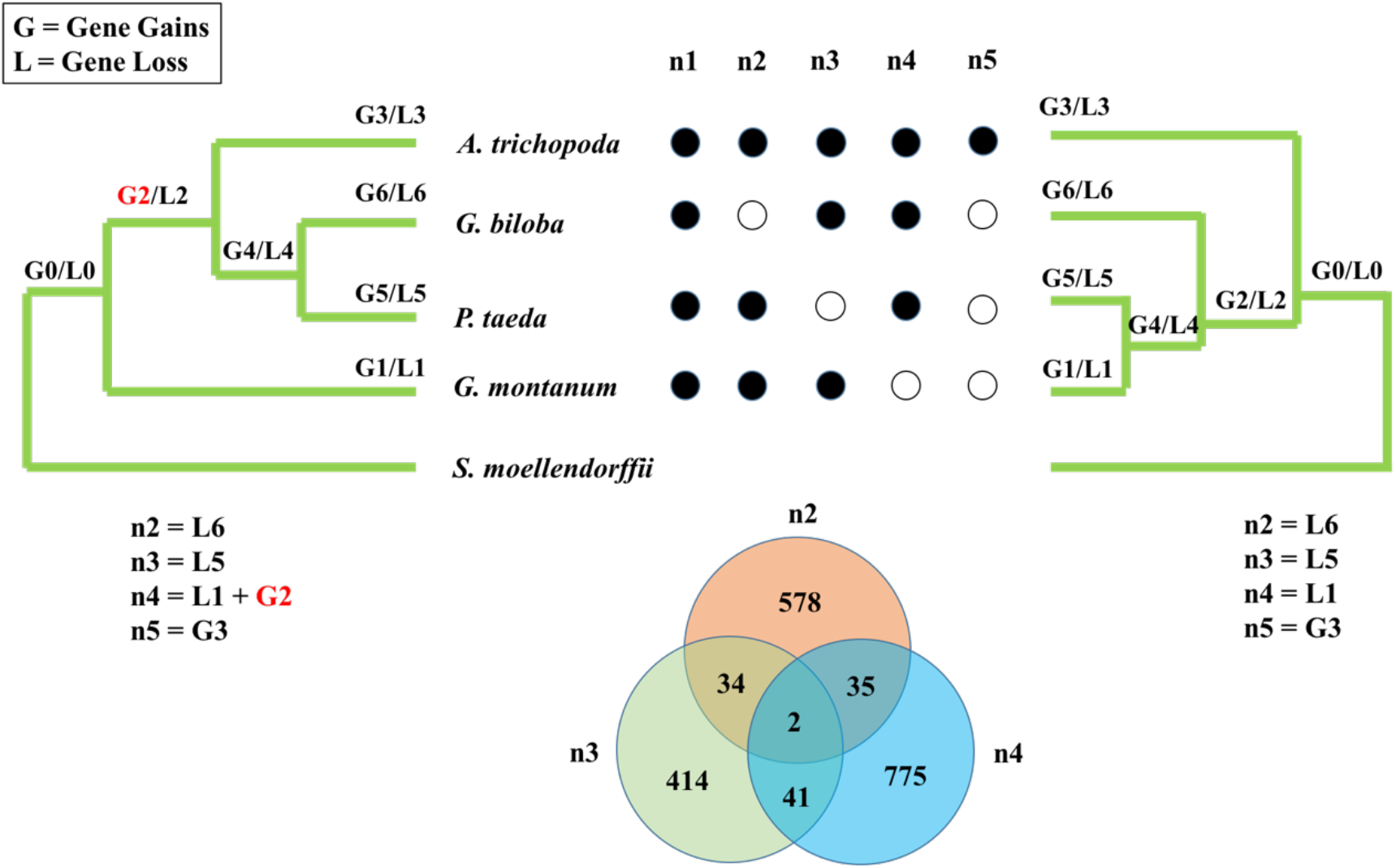
Interpretation of distinct Genic signatures for gnetophytes based on gene gains/loss mechanism with ‘Seed plants sister’(right) hypothesis and ‘closed to conifers’ hypothesis. More gene loss detected in *G. montanum* and ‘n2’ genes harboring less identity is due the G2 part of genes (genes gained by the MRCA of other seed plants). The right topology tree can not explain the distinct patterns found in *G. montanum*.

For additional, we examined in detail some exemplar multigene families of the “n4” set using phylogenetic approaches and observed two families that are consistent with the ‘Seed plants sister’ hypothesis. For example, from the germin-like protein family the orthologue GLP7 is missing from *G. montanum* and is broadly shared by non-gnetophytes gymnosperms and angiosperms (see Supplementary Materials online and Supplementary Fig. 1). In the same way, sub-clades of the Phenylalanine Ammonia Lyase gene family are found to be only shared by non-gnetophytes gymnosperms and angiosperms (Supplementary Fig. 2, Supplementary Table 4; Bagal et al., 2012).

## Discussion

We argue here that the hypotheses ‘gnetophytes are sister to all other seed plants’ can not be ruled out based only on the phylogenetic trees inferred from nuclear loci. Assuming so, there remains the potential for profound shifts in our understanding of the origin and divergence of many genetic, genomic, biochemical, metabolic and morphological characters in seed plant evolution. For example, the two paralogues GgWOXX and GgWOXY should be consider to have been lost in the their MRCA of other seed plants after the split with gnetophytes (Wan et al., 2018). And if gymnosperms may not be monophyletic, then many characters used to be considered to be derived in gnetophytes (Wan et al., 2018) may in fact be ancestral characters (Fig. 5, e.g. intron structures, lack of WGD, pfam domains).

**Figure 5.**
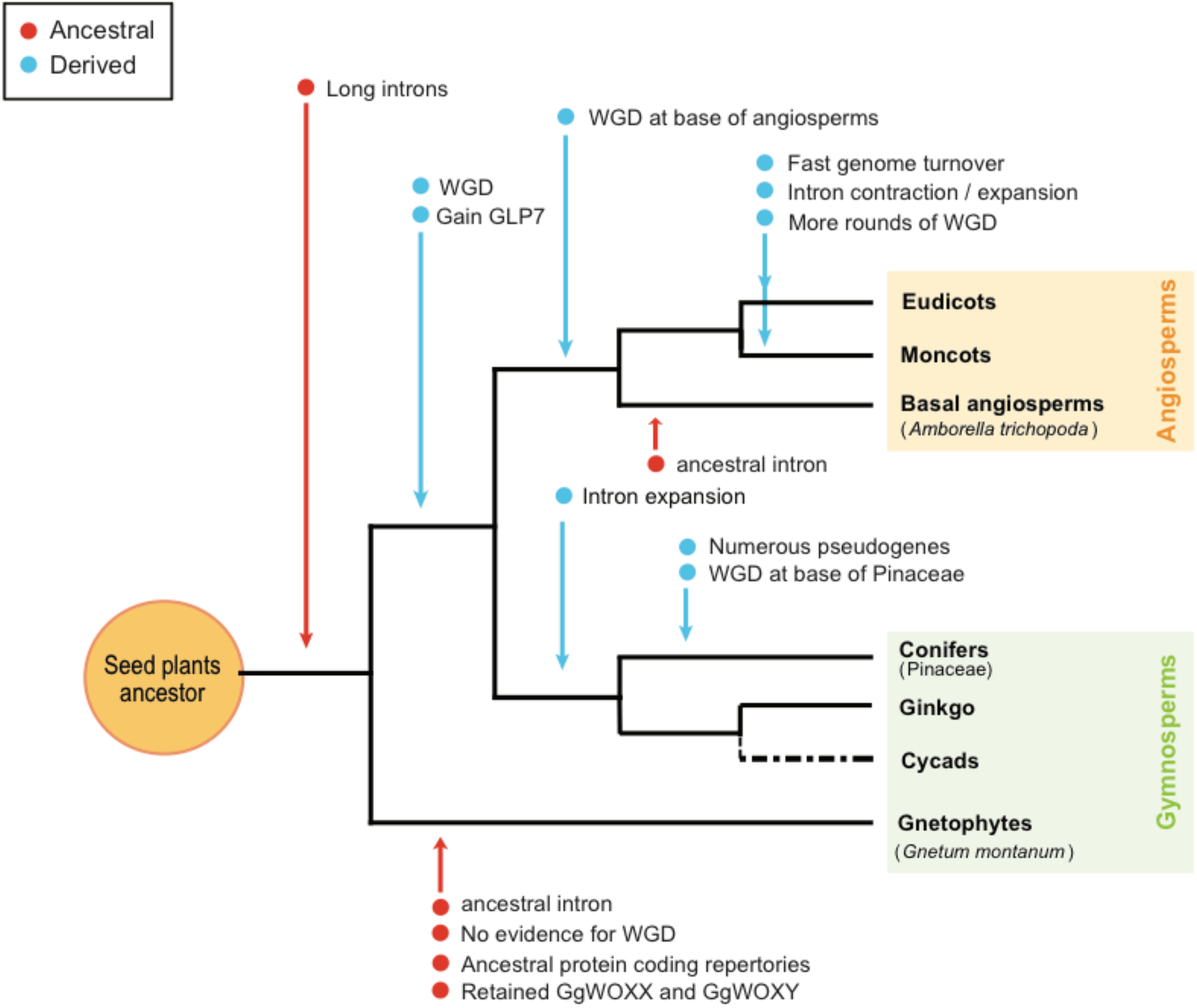
Reinterpretation of genome evolution patterns across seed plants based on the hypotheses gnetophytes are sister to all other seed plants.

## ACKNOWLEDGEMENTS

Thanks to Prof. Antonis Rokas, who provide guidance and formula on gene-wise log-likelihood scores (GLS), and also to Dr. Yanting Hu for her kindly help on Cycads transcriptome data transfer. We also thank Prof. Andrew R. Leitch, Ilia J. Leitch, Yves Van de Peer, Laura Kelly, Honglei Li, Yang Liu, Min Yang, Qingfeng Wang, Tao Wan, Jinbo Zhang, and Ji Li for the useful comments on the manuscript.

## AUTHOR CONTRIBUTIONS

Yinzhi Zhang conceived the study and led the manuscript preparation. Yinzhi Zhang worked on the data matrix construction; Zhiming Liu mostly contributed to the phylogenetic analyses of PAL, GLP families. Yinzhi Zhang mostly contributed to the analyses of gene family divergent pattern comparison.

## ADDITIONAL INFORMATON

Supplementary information including materials, methods, figures and tables are available at *BioRxiv* Online.

