## Supplementary Figures for "Genic evidence that gnetophytes are sister to all other seed plants"

### Supplementary materials and methods

#### Phylogenetic reconstruction of Germin-like proteins (GLP) and Phenylalanine Ammonia Lyase (PAL) gene families

The "n4" gene set (= 853 genes considered absent in *G. montanum*, supplementary table S9) contained members of the GLP family, a large multigene family of ubiquitous plant glycoproteins involved in seedling germination, and abiotic (e.g. drought, aluminium toxicity) and biotic stress (Membre et al. 2000; Bernier and Berna 2001). For the identification of GLPs, we retrieved the conserved domains of GLP (PF00190) from (i) the pfam database (http://pfam.xfam.org/) using eight representative plant genomes (*S. moellendorffii*, *P. taeda*, *P. abies*, *G. biloba*, *A. trichopoda*, *A. thaliana*, *G. montanum*, and *E. equisetina*) (ii) *O. sativa*'s GLP protein sequences. These were aligned using HMMER (v3.1b2, http://hmmer.org) and only proteins having the corresponding conserved domains were retained. We then used MUSCLE (Edgar 2004) (v3.8.31; http://www.drive5.com/muscle) with default parameters for multiple sequence alignments and we constructed the phylogenetic trees using Maximum Likelihood (ML) methods with RAxML (Stamatakis, et al. 2005) (v8.0.19, parameter: '-m PROTGAMMAAUTO -p 12345 -x 12345 -# 20 -f ad -T 20'). The clades of the phylogenetic trees were mainly labeled using the genes from *A. thaliana* on the TAIR databases (https://www.arabidopsis.org/). The result showed several distinct clades that were specific to certain taxonomic groups (e.g. lycophytes, angiosperms), and some (GLP1/3) that were ubiquitous to all seed plants analysed. Critically, however, one well-supported clade (GLP7), previously used to trace GLP family diversity across seed plants (Carter and Thornburg 1999) was found in all seed plants except gnetophytes (Supplementary Fig. 1).

PAL gene family is another example within the 'n4' gene set. Previous work has identified many distinct PAL genes in gymnosperms, and they have been shown to be much more diverse compared with angiosperms (Bagal et al. 2012). For the identification of PAL, we retrieved (i) the conserved domains (PF00221) from the pfam database (http://pfam.xfam.org/) from seven representative plant genomes: *P. taeda*, *P. abies*, *G. biloba*, *A. trichopoda*, *A. thaliana*, *G. montanum*, and *E. equisetina*; (ii) PAL protein sequences obtained from a previous study (Ritter and Schulz 2004; Bagal, et al. 2012), and; (iii) PAL sequences of *S. moellendorffii* from the NCBI website (https://www.ncbi.nlm.nih.gov). We aligned these sequences using a HMMER (v3.1b2, http://hmmer.org) search and only proteins having the corresponding conserved domains were retained. We then used MUSCLE (Edgar 2004) (v3.8.31; http://www.drive5.com/muscle) with default parameters for multiple sequence alignments and we constructed phylogenetic trees using ML methods with RAxML (Stamatakis, et al. 2005) (v8.0.19, parameter set as: '-m PROTGAMMAAUTO -p 12345 -x 12345 -# 100 -f ad -T 20'). The clades of the phylogenetic trees were mainly labeled using the gene from *A. thaliana* on the TAIR databases (https://www.arabidopsis.org/). Result showed that no gnetophyte sequences were found in the clades (group 3 and group 4) which are likely to have occurred with the divergence of gymnosperms and angiosperms (Bagal, et al. 2012) (Supplementary Fig. 2).

**Supplementary Table 1-5. Information of orthoMCL protein genes in the five categorized sets from the sampled seed plant species.**

(See separate Excel file)

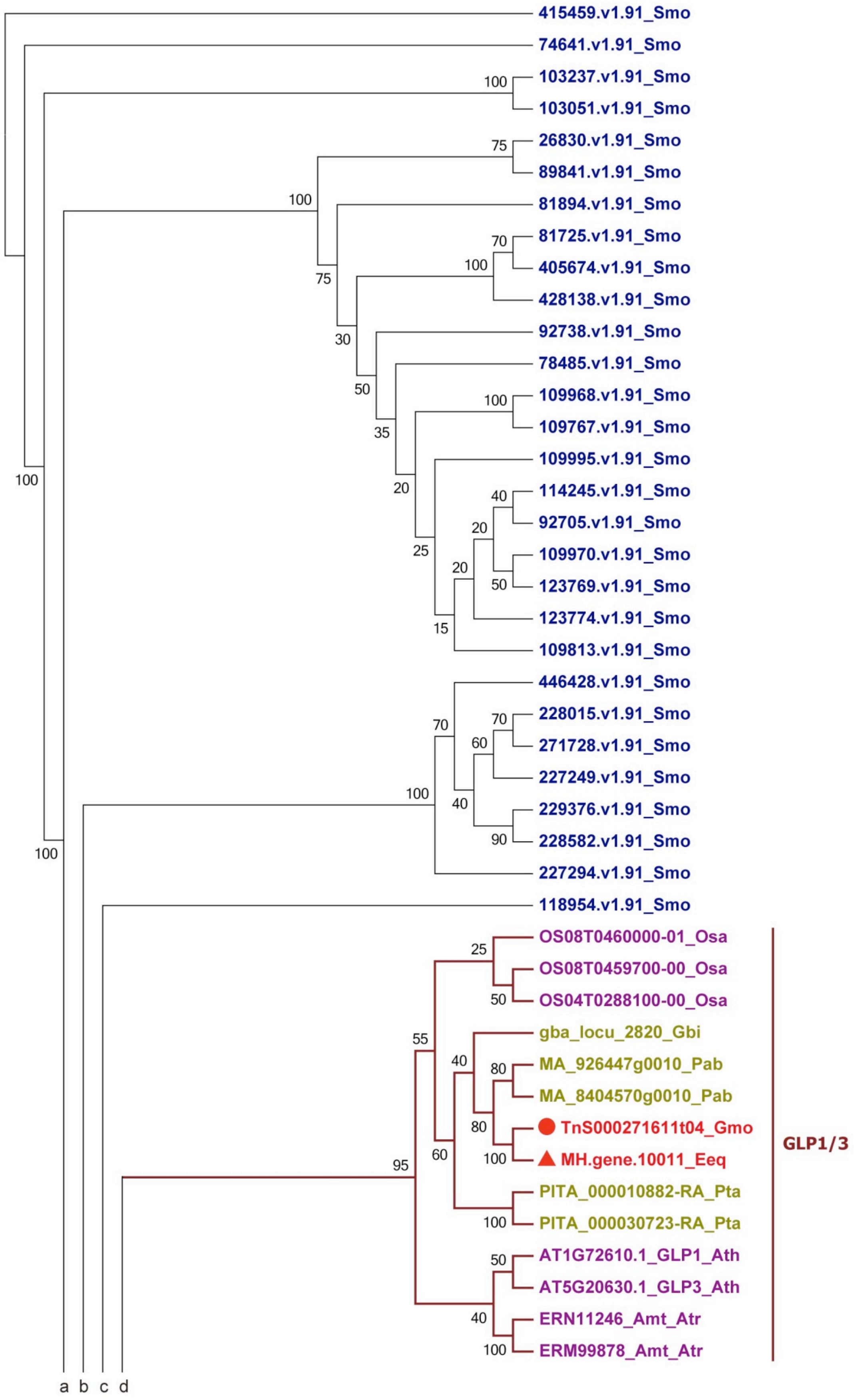
415459.v1.91_Smo
74641.v1.91_Smo
103237.v1.91_Smo
103051.v1.91_Smo
26830.v1.91_Smo
89841.v1.91_Smo
81894.v1.91_Smo
81725.v1.91_Smo
405674.v1.91_Smo
428138.v1.91_Smo
92738.v1.91_Smo
78485.v1.91_Smo
109968.v1.91_Smo
109767.v1.91_Smo
109995.v1.91_Smo
114245.v1.91_Smo
92705.v1.91_Smo
109970.v1.91_Smo
123769.v1.91_Smo
123774.v1.91_Smo
109813.v1.91_Smo
446428.v1.91_Smo
228015.v1.91_Smo
271728.v1.91_Smo
227249.v1.91_Smo
229376.v1.91_Smo
228582.v1.91_Smo
227294.v1.91_Smo
118954.v1.91_Smo
OS08T0460000-01_Osa
OS08T0459700-00_Osa
OS04T0288100-00_Osa
gba_locu_2820_Gbi
MA_926447g0010_Pab
MA_8404570g0010_Pab
TnS000271611t04_Gmo
MH.gene.10011_Eeq
PITA_000010882-RA_Pta
PITA_000030723-RA_Pta
AT1G72610.1_GLP1_Ath
AT5G20630.1_GLP3_Ath
ERN11246_Amt_Atr
ERM99878_Amt_Atr
GLP1/3
a b c d

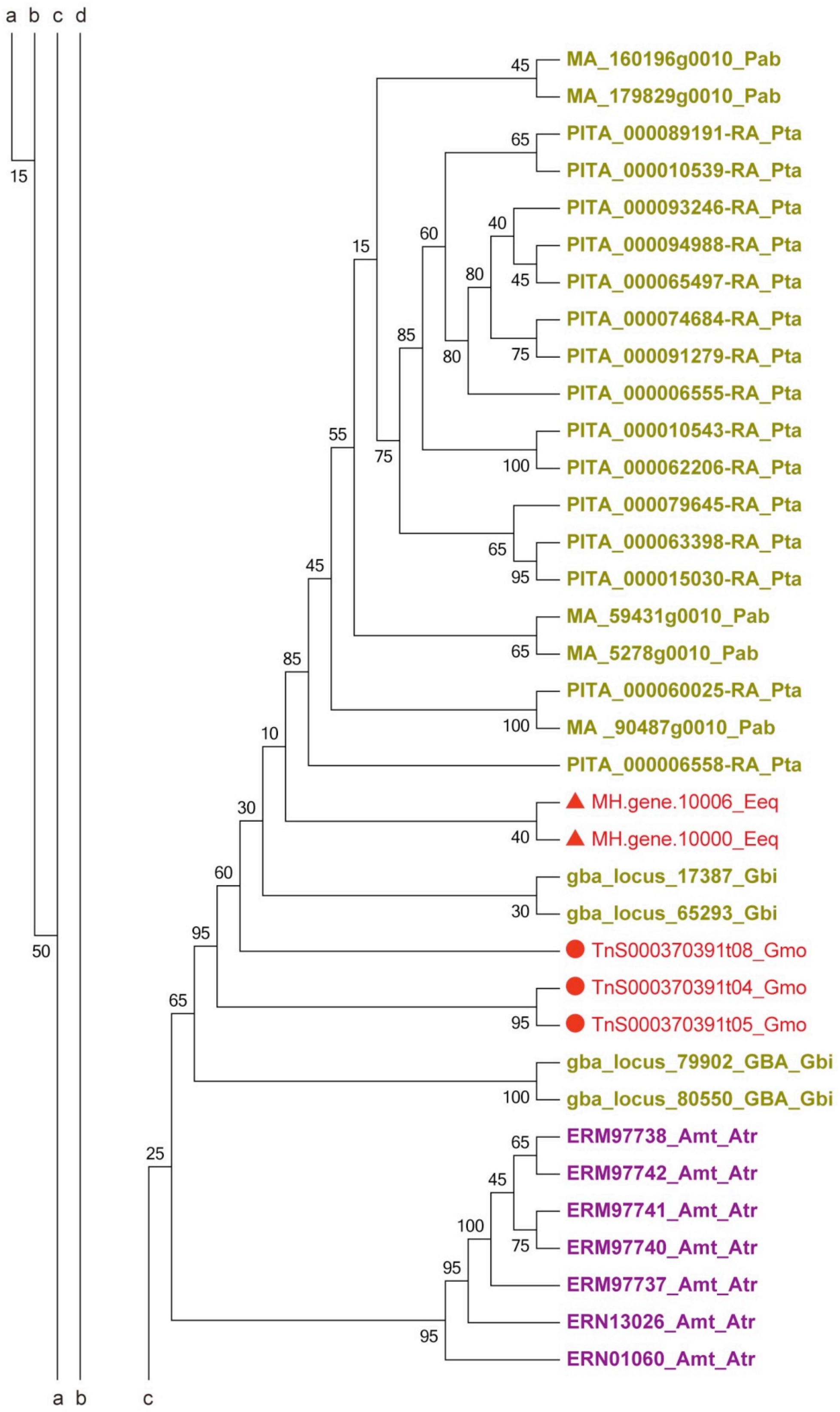

a b c d
MA_160196g0010_Pab
MA_179829g0010_Pab
PITA_000089191-RA_Pta
PITA_000010539-RA_Pta
PITA_000093246-RA_Pta
PITA_000094988-RA_Pta
PITA_000065497-RA_Pta
PITA_000074684-RA_Pta
PITA_000091279-RA_Pta
PITA_000006555-RA_Pta
PITA_000010543-RA_Pta
PITA_000062206-RA_Pta
PITA_000079645-RA_Pta
PITA_000063398-RA_Pta
PITA_000015030-RA_Pta
MA_59431g0010_Pab
MA_5278g0010_Pab
PITA_000060025-RA_Pta
MA _90487g0010_Pab
PITA_000006558-RA_Pta
MH.gene.10006_Eeq
MH.gene.10000_Eeq
gba_locus_17387_Gbi
gba_locus_65293_Gbi
TnS000370391t08_Gmo
TnS000370391t04_Gmo
TnS000370391t05_Gmo
gba_locus_79902_GBA_Gbi
gba_locus_80550_GBA_Gbi
ERM97738_Amt_Atr
ERM97742_Amt_Atr
ERM97741_Amt_Atr
ERM97740_Amt_Atr
ERM97737_Amt_Atr
ERN13026_Amt_Atr
ERN01060_Amt_Atr
a b c

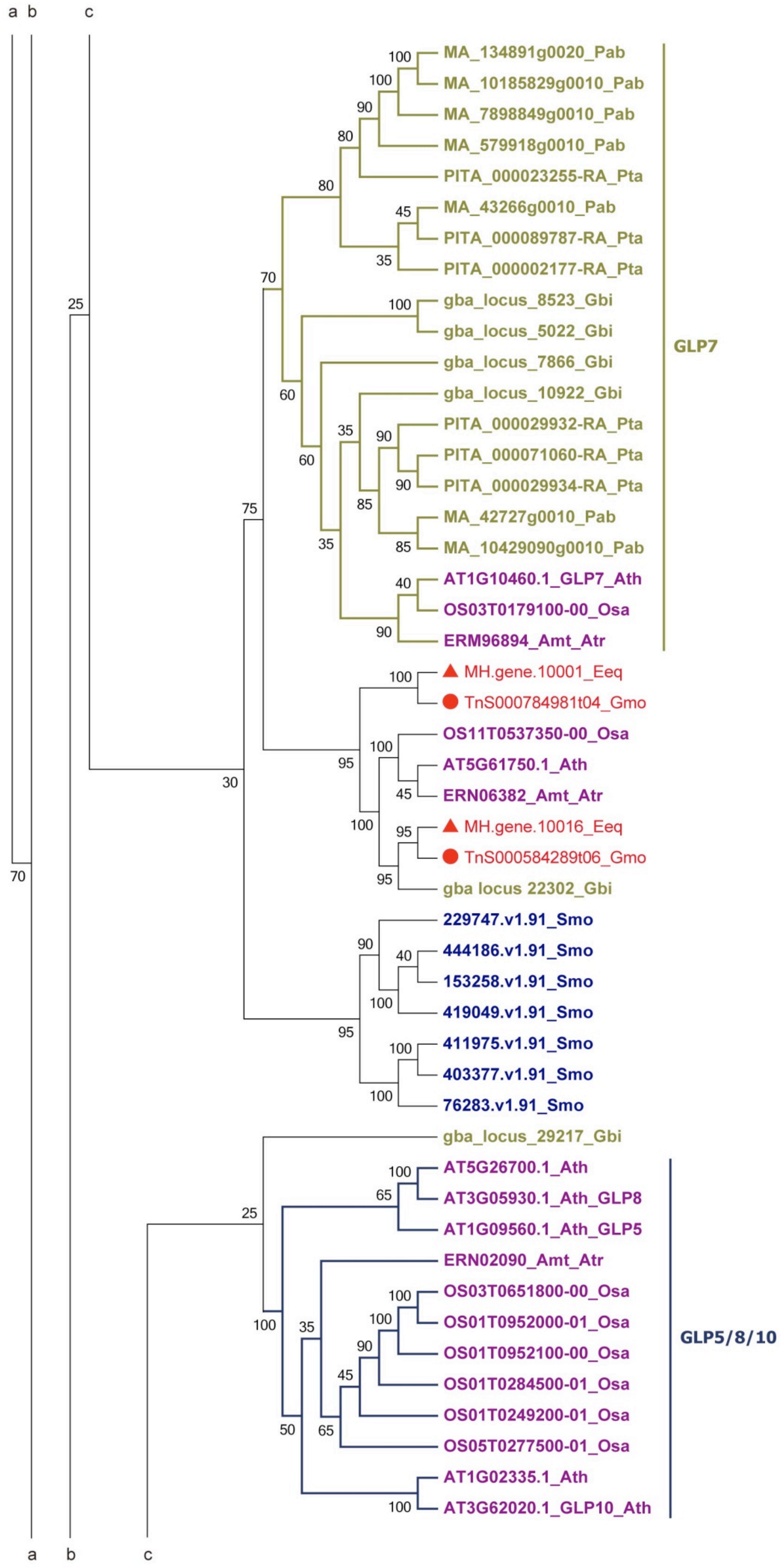

a b c
MA_134891g0020_Pab
MA_10185829g0010_Pab
MA_7898849g0010_Pab
MA_579918g0010_Pab
PITA_000023255-RA_Pta
MA_43266g0010_Pab
PITA_000089787-RA_Pta
PITA_000002177-RA_Pta
gba_locus_8523_Gbi
gba_locus_5022_Gbi
gba_locus_7866_Gbi
gba_locus_10922_Gbi
PITA_000029932-RA_Pta
PITA_000071060-RA_Pta
PITA_000029934-RA_Pta
MA_42727g0010_Pab
MA_10429090g0010_Pab
AT1G10460.1_GLP7_Ath
OS03T0179100-00_Osa
ERM96894_Amt_Atr
GLP7
MH.gene.10001_Eeq
TnS000784981t04_Gmo
OS11T0537350-00_Osa
AT5G61750.1_Ath
ERN06382_Amt_Atr
MH.gene.10016_Eeq
TnS000584289t06_Gmo
gba locus 22302_Gbi
229747.v1.91_Smo
444186.v1.91_Smo
153258.v1.91_Smo
419049.v1.91_Smo
411975.v1.91_Smo
403377.v1.91_Smo
76283.v1.91_Smo
gba_locus_29217_Gbi
AT5G26700.1_Ath
AT3G05930.1_Ath_GLP8
AT1G09560.1_Ath_GLP5
ERN02090_Amt_Atr
OS03T0651800-00_Osa
OS01T0952000-01_Osa
OS01T0952100-00_Osa
OS01T0284500-01_Osa
OS01T0249200-01_Osa
OS05T0277500-01_Osa
AT1G02335.1_Ath
AT3G62020.1_GLP10_Ath
GLP5/8/10
a b c

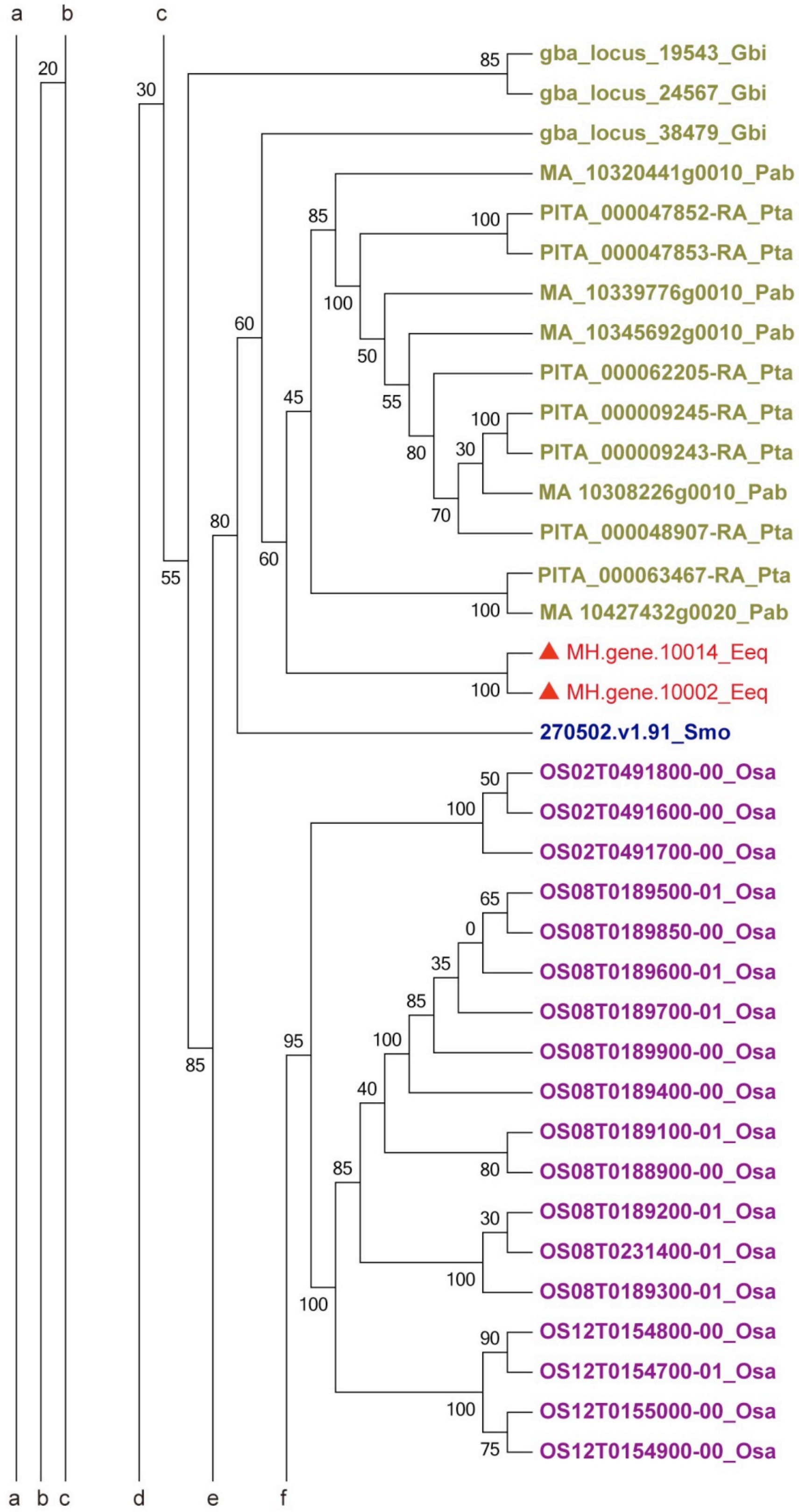
a
b
c
gba_locus_19543_Gbi
gba_locus_24567_Gbi
gba_locus_38479_Gbi
MA_10320441g0010_Pab
PITA_000047852-RA_Pta
PITA_000047853-RA_Pta
MA_10339776g0010_Pab
MA_10345692g0010_Pab
PITA_000062205-RA_Pta
PITA_000009245-RA_Pta
PITA_000009243-RA_Pta
MA 10308226g0010_Pab
PITA_000048907-RA_Pta
PITA_000063467-RA_Pta
MA 10427432g0020_Pab
MH.gene.10014_Eeq
MH.gene.10002_Eeq
270502.v1.91_Smo
OS02T0491800-00_Osa
OS02T0491600-00_Osa
OS02T0491700-00_Osa
OS08T0189500-01_Osa
OS08T0189850-00_Osa
OS08T0189600-01_Osa
OS08T0189700-01_Osa
OS08T0189900-00_Osa
OS08T0189400-00_Osa
OS08T0189100-01_Osa
OS08T0188900-00_Osa
OS08T0189200-01_Osa
OS08T0231400-01_Osa
OS08T0189300-01_Osa
OS12T0154800-00_Osa
OS12T0154700-01_Osa
OS12T0155000-00_Osa
OS12T0154900-00_Osa
a b c d e f

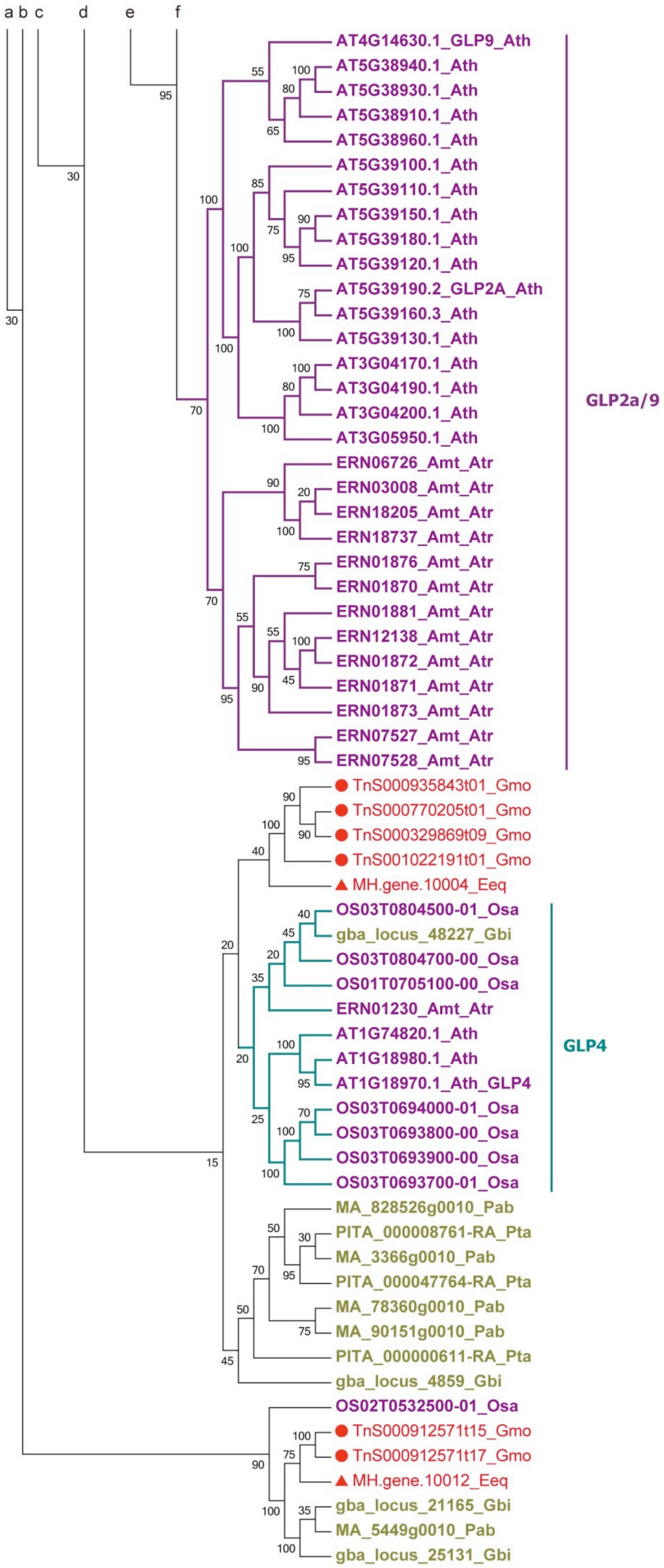

a b c d e f
AT4G14630.1_GLP9_Ath
AT5G38940.1_Ath
AT5G38930.1_Ath
AT5G38910.1_Ath
AT5G38960.1_Ath
AT5G39100.1_Ath
AT5G39110.1_Ath
AT5G39150.1_Ath
AT5G39180.1_Ath
AT5G39120.1_Ath
AT5G39190.2_GLP2A_Ath
AT5G39160.3_Ath
AT5G39130.1_Ath
AT3G04170.1_Ath
AT3G04190.1_Ath
AT3G04200.1_Ath
AT3G05950.1_Ath
ERN06726_Amt_Atr
ERN03008_Amt_Atr
ERN18205_Amt_Atr
ERN18737_Amt_Atr
ERN01876_Amt_Atr
ERN01870_Amt_Atr
ERN01881_Amt_Atr
ERN12138_Amt_Atr
ERN01872_Amt_Atr
ERN01871_Amt_Atr
ERN01873_Amt_Atr
ERN07527_Amt_Atr
ERN07528_Amt_Atr
GLP2a/9
TnS000935843t01_Gmo
TnS000770205t01_Gmo
TnS000329869t09_Gmo
TnS001022191t01_Gmo
MH.gene.10004_Eeq
OS03T0804500-01_Osa
gba_locus_48227_Gbi
OS03T0804700-00_Osa
OS01T0705100-00_Osa
ERN01230_Amt_Atr
AT1G74820.1_Ath
AT1G18980.1_Ath
AT1G18970.1_Ath_GLP4
OS03T0694000-01_Osa
OS03T0693800-00_Osa
OS03T0693900-00_Osa
OS03T0693700-01_Osa
GLP4
MA_828526g0010_Pab
PITA_000008761-RA_Pta
MA_3366g0010_Pab
PITA_000047764-RA_Pta
MA_78360g0010_Pab
MA_90151g0010_Pab
PITA_000000611-RA_Pta
gba_locus_4859_Gbi
OS02T0532500-01_Osa
TnS000912571t15_Gmo
TnS000912571t17_Gmo
MH.gene.10012_Eeq
gba_locus_21165_Gbi
MA_5449g0010_Pab
gba_locus_25131_Gbi

**Supplementary Fig. 1. Phylogenetic analysis of the Germin-Like Protein (GLP) gene family.** Abbreviations of species name after the sequences listed as: *S. moellendorffii* (Smo), *A. trichopoda* (Atr), *A. thaliana* (Ath), *O. sativa* (Osa), *G. montanum* (Gmo), *E. equisetina* (Eeq), *P. abies* (Pab), *P. taeda* (Pta), *G. biloba* (Gbi). Red circles and triangles indicate members from *G. montanum* and *E. equisetina*, respectively. The analysis resolves several clades, including: (i) GLP1/3 is a well-supported clade (95%) of sequences that are multicopy in angiosperms and conifers and single copy in gnetophytes and *G. biloba*. Interpretation - expansion of sequences occurred after gnetophytes split from the rest of seed plants. (ii) GLP7 is a moderately-supported (70%) clade comprising sequences that are multicopy in non-gnetophyte gymnosperms and single copy in angiosperms. Interpretation - GLP7 diverged and amplified after gnetophytes split from the rest of seed plants. (iii) GLP5/8/10 is a well - supported (100%) clade and GLP2a/9 moderately supported clade (70%) comprising only angiosperms. Interpretation - diverged and proliferated within angiosperms. (iv) Well-supported clade (90%) containing sequences from angiosperms, *G. biloba*, conifers and gnetophytes. Interpretation - sequences diverged with ancestral seed plants. (v) A well-supported (100%) clade of *G. montanum* sequences. Interpretation - sequences diverged with *G. montanum* (gnetophytes) (vi) A well-supported clade (80%) of gnetophytes, *G. biloba*, conifer sequences and S. *moellendorffii* sequences. Interpretation - sequence in ancestral land plants lost in angiosperms. (vii) A well-supported (95%) clade including paralogues clustering to species in gnetophytes, *G. biloba* and conifer sequences. Interpretation - Ancestral seed plant sequence lost in angiosperms (or gained in common ancestor to all gymnosperms) and amplified along each lineage independently. (ix) A well-supported clade (96%) including sequences in gnetophytes, angiosperms, and *G. biloba*. Interpretation - sequence in ancestral seed plants lost in conifers. None of these sequences / clades support an interpretation of gnetophytes sister to, or embedded within conifers.

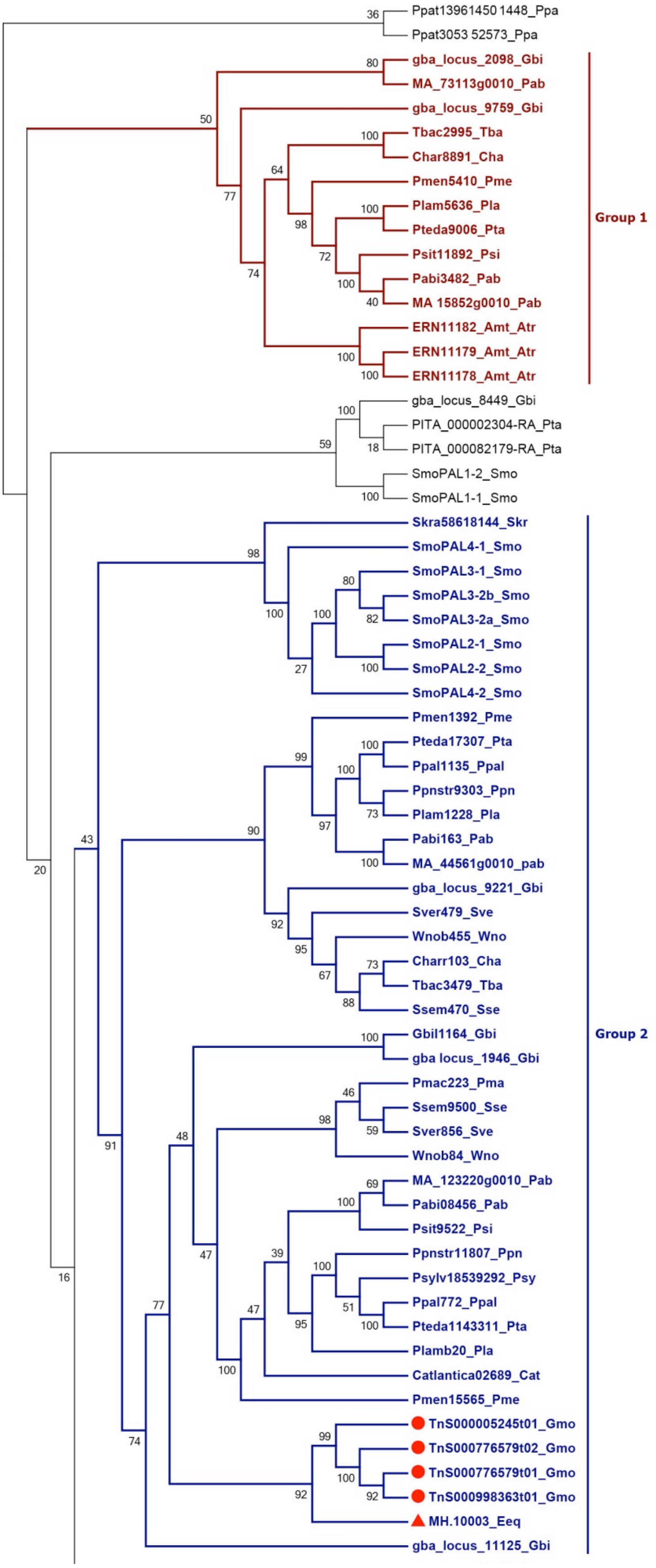

Ppat13961450 1448_Ppa
Ppat3053 52573_Ppa
gba_locus_2098_Gbi
MA_73113g0010_Pab
gba_locus_9759_Gbi
Tbac2995_Tba
Char8891_Cha
Pmen5410_Pme
Plam5636_Pla
Pteda9006_Pta
Psit11892_Psi
Pabi3482_Pab
MA 15852g0010_Pab
ERN11182_Amt_Atr
ERN11179_Amt_Atr
ERN11178_Amt_Atr
Group 1
gba_locus_8449_Gbi
PITA_000002304-RA_Pta
PITA_000082179-RA_Pta
SmoPAL1-2_Smo
SmoPAL1-1_Smo
Skra58618144_Skr
SmoPAL4-1_Smo
SmoPAL3-1_Smo
SmoPAL3-2b_Smo
SmoPAL3-2a_Smo
SmoPAL2-1_Smo
SmoPAL2-2_Smo
SmoPAL4-2_Smo
Pmen1392_Pme
Pteda17307_Pta
Ppal1135_Ppal
Ppnstr9303_Ppn
Plam1228_Pla
Pabi163_Pab
MA_44561g0010_pab
gba_locus_9221_Gbi
Sver479_Sve
Wnob455_Wno
Charr103_Cha
Tbac3479_Tba
Ssem470_Sse
Gbil1164_Gbi
gba locus_1946_Gbi
Pmac223_Pma
Ssem9500_Sse
Sver856_Sve
Wnob84_Wno
MA_123220g0010_Pab
Pabi08456_Pab
Psit9522_Psi
Ppnstr11807_Ppn
Psylv18539292_Psy
Ppal772_Ppal
Pteda1143311_Pta
Plamb20_Pla
Catlantica02689_Cat
Pmen15565_Pme
TnS000005245t01_Gmo
TnS000776579t02_Gmo
TnS000776579t01_Gmo
TnS000998363t01_Gmo
MH.10003_Eeq
gba_locus_11125_Gbi
Group 2

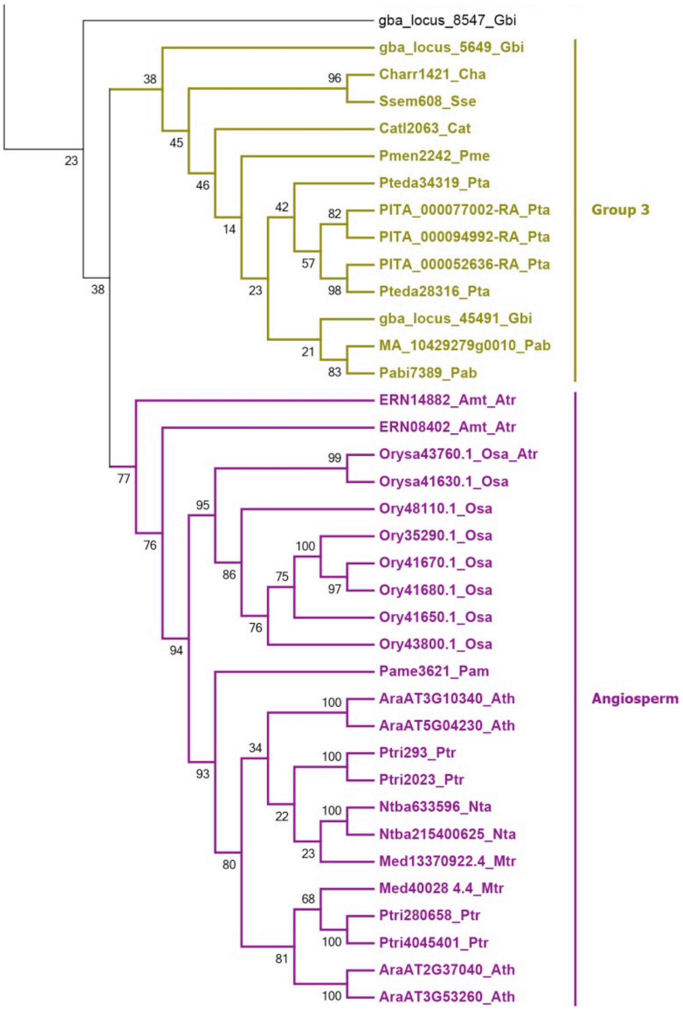


**Supplementary Fig. 2. Phylogenetic analysis of the PAL gene family.** Abbreviation of species name after the sequences listed as: *P. patens* (Ppa), *S. moellendorffii* (Smo), *Selaginella kraussiana* (Skr), *G. montanum* (Gmo), *E .equisetina* (Eeq), *P. abies* (Pab), *P. taeda* (Pta), *G. biloba* (Gbi), *Cupressus atlantica* (Cat), *Cephalotaxus harringtonia* (Cha), *Picea sitchensis* (Psi), *Pinus lambertiana* (Pla), *Pseudotsuga menziesii* (Pme), *Pinus palustris* (Ppal), *Pinus pinaster* (Ppi), *Pinus sylvestris* (Psy), *Pinus taeda* (Pta), *Podocarpus macrophyllus* (Pma), *Sciadopitys verticillata* (Sve), *Sequoia sempervirens* (Sse), *Taxus baccata* (Tba), *Wollemia nobilis* (Wno), *A. trichopoda* (Atr), *Medicago truncatula* (Mtr), *Nicotiana tabacum* (Nta), *Persea americana* (Pam), *P. trichocarpa* (Ptr), *O. sativa* (Osa), *A. thaliana* (Ath). Red circles and triangles indicate members from *G. montanum* and *E. equisetina*, respectively. The distribution of PAL family members show (i) Multiple paralogues and orthologues within an angiosperm clade. Interpretation - sequences diverged after angiosperms split from the rest of seed plants. (ii) Multiple paralogues and orthologues from conifers and *G. biloba* in separate clades of varying support, some with unclear relationships to orthologues from non-seed plants. Interpretation - divergence and amplification of genes in lineages leading to conifers and *G. biloba*. (iii) Multiple paralogues forming a clade of gnetophyte sequences, with unresolved relationships to sequences from conifers, *G.*

*biloba* and non-seed plants. Interpretation - a gnetophyte lineage-specific amplification of a subset of the diversity found in conifers and *G. biloba*. The absence of gnetophyte orthologues associated with the remaining conifer / *G.biloba* diversity might suggest the early divergence of gnetophytes from the rest of gymnosperms.
